# Functional Maturation of Human iPSC-based Cardiac Microphysiological Systems with Tunable Electroconductive Decellularized Extracellular Matrices

**DOI:** 10.1101/786657

**Authors:** Jonathan H. Tsui, Andrea Leonard, Nathan D. Camp, Joseph T. Long, Zeid Y. Nawas, Rakchanok Chavanachat, Jong Seob Choi, Alejandro Wolf-Yadlin, Charles E. Murry, Nathan J. Sniadecki, Deok-Ho Kim

## Abstract

Human induced pluripotent stem cell-derived cardiomyocytes (hiPSC-CMs) offer tremendous potential for use in engineering human tissues for regenerative therapy and drug screening. However, differentiated cardiomyocytes are phenotypically immature, reducing assay reliability when translating *in vitro* results to clinical studies and precluding hiPSC-derived cardiac tissues from therapeutic use *in vivo*. To address this, we have developed hybrid hydrogels comprised of decellularized porcine myocardial extracellular matrix (dECM) and reduced graphene oxide (rGO) to provide a more instructive microenvironment for proper cellular and tissue development. A tissue-specific protein profile was preserved post-decellularization, and through the modulation of rGO content and degree of reduction, the mechanical and electrical properties of the hydrogels could be tuned. Engineered heart tissues (EHTs) generated using dECM-rGO hydrogel scaffolds and hiPSC-derived cardiomyocytes exhibited significantly increased twitch forces at 14 days of culture and had increased the expression of genes that regulate contractile function. Similar improvements in various aspects of electrophysiological function, such as calcium-handling, action potential duration, and conduction velocity, were also induced by the hybrid biomaterial. We also demonstrate that dECM-rGO hydrogels can be used as a bioink to print cardiac tissues in a high-throughput manner, and these tissues were utilized to assess the proarrhythmic potential of cisapride. Action potential prolongation and beat interval irregularities was observed in dECM-rGO tissues at clinical doses of cisapride, indicating that the enhanced maturation of these tissues corresponded well with a capability to produce physiologically relevant drug responses.

---

Cardiovascular disease (CVD) remains a leading cause of death for both men and women worldwide, with the economic costs of healthcare related to treatment or quality of life improvements forecasted to exceed $1 trillion per year in the United States alone by 2030^1, 2^. With the healthcare burden of CVD increasing every year, considerable research is devoted to engineering cardiac tissues to better understand the underlying disease pathologies, to screen potential pharmacological treatments, and to serve as implantable therapies. The non-proliferative nature of terminally differentiated cardiomyocytes has served as a bottleneck for such work, as obtaining sufficient numbers of cells for studies is ethically and logistically difficult. The advent of human induced pluripotent stem cells (hiPSCs) has somewhat ameliorated this issue since not only are hiPSCs readily obtainable and scalable in culture, hiPSC lines can now be generated from patients with specific disease phenotypes^3, 4^. However, cardiomyocytes differentiated from hiPSCs are phenotypically immature and behave more like cells at a neonatal, rather than adult, stage of development^5^. This reduces reliability when translating *in vitro* assay results to clinical studies, as immature cells do not accurately reproduce the pharmacological responses and disease progression of the native myocardium.

To address this issue with cardiomyocyte maturity, various strategies have been studied in the field. For examples, studies have demonstrated that preconditioning of engineered cardiac tissues with specific biochemical, mechanical, and electrical stimulation regimes elicited beneficial effects on overall tissue function and maturation^6–8^. However, these techniques can be limited by their requirement for the integration of complex hardware and software, thereby resulting in potential difficulties with platform adaptability and in increasing experimental throughput. Alternatively, improved maturity can be potentially achieved by exploring attempts to recapitulate the myocardial extracellular matrix (ECM) that defines the architecture, signaling, and biomechanics of cardiomyocytes *in vivo*^9, 10^, and by utilizing engineered ECMs, mature tissues could be generated across a variety of existing platforms. Recent advances in developing three-dimensional (3D) matrices have been able to better reproduce this cell niche; however, overall cardiomyocyte and engineered cardiac tissue maturity is still at suboptimal levels due to limitations with the scaffolds and platforms that are currently available. Naturally-derived polymers typically used to engineer 3D tissues, such as fibrin and collagen, offer good biocompatibility and biochemical similarity to *in vivo* systems. In particular, decellularized extracellular matrices (dECM) are able to largely preserve the tissue-specific biochemical makeup of the tissues that they are derived from^11^. The poor mechanical and electrical properties of these polymers, however, often hampers their ability to replicate the tissue microenvironment. Synthetic materials on the other hand, offer better control over their physical and chemical characteristics, but their bioactivity is often non-existent without further modification. Perhaps the most interesting exceptions are highly-electroconductive materials, such as carbon nanotubes^12, 13^, graphene derivatives^14, 15^, and gold^16, 17^, that have been found to impart beneficial effects on the growth and maturation of a variety of cell types and have recently gained increasing prominence in the field of tissue engineering.

Here, we report on the development of hybrid hydrogels that are comprised of reduced graphene oxide (rGO) dispersed within decellularized myocardial extracellular matrices (dECM). By leveraging the advantages offered by rGO with regards to controlling properties such as stiffness and electroconductivity while maintaining the bioactivity inherent with dECM, this new material can advance the capability to engineer 3D human cardiac tissues that are more physiologically-representative of the native myocardium.

## Results

### Synthesis and characterization of dECM-rGO composite hydrogels with tunable mechanical and electrical properties

The two primary components of the dECM-rGO composite hydrogels used in this study were prepared by means of parallel processes (**Fig. 1a**). Left ventricular myocardium from freshly-harvested porcine hearts were decellularized using an established method that has been previously demonstrated to produce dECM in which the majority of cellular components have been removed, while still preserving the collagens and glycosaminoglycans (GAGs) originally present in the tissue^18^. As one of the stated benefits of utilizing dECM as a scaffold material is its recapitulation of the native tissue microenvironment, further analysis of its biochemical makeup post-decellularization was conducted using liquid chromatography tandem mass spectrometry (LC-MS/MS). Several produced batches of dECM were analyzed for key matrix proteins: collagens I – IV, fibronectin, and laminin (**Fig. 1b**). While batch-to-batch variation in the quantities of these proteins did exist, the overall content and relative ratios were well maintained (**Fig. 1c**), with collagen I the most abundant protein type found in the myocardial dECM. The next most abundant proteins found were collagen III, fibronectin, and laminin. This ECM protein composition is similar to what has been found in adult rat hearts^19^, indicating that the dECM produced maintains a developmental stage-appropriate tissue-specificity in regards to biochemical signaling.

**Figure 1.**
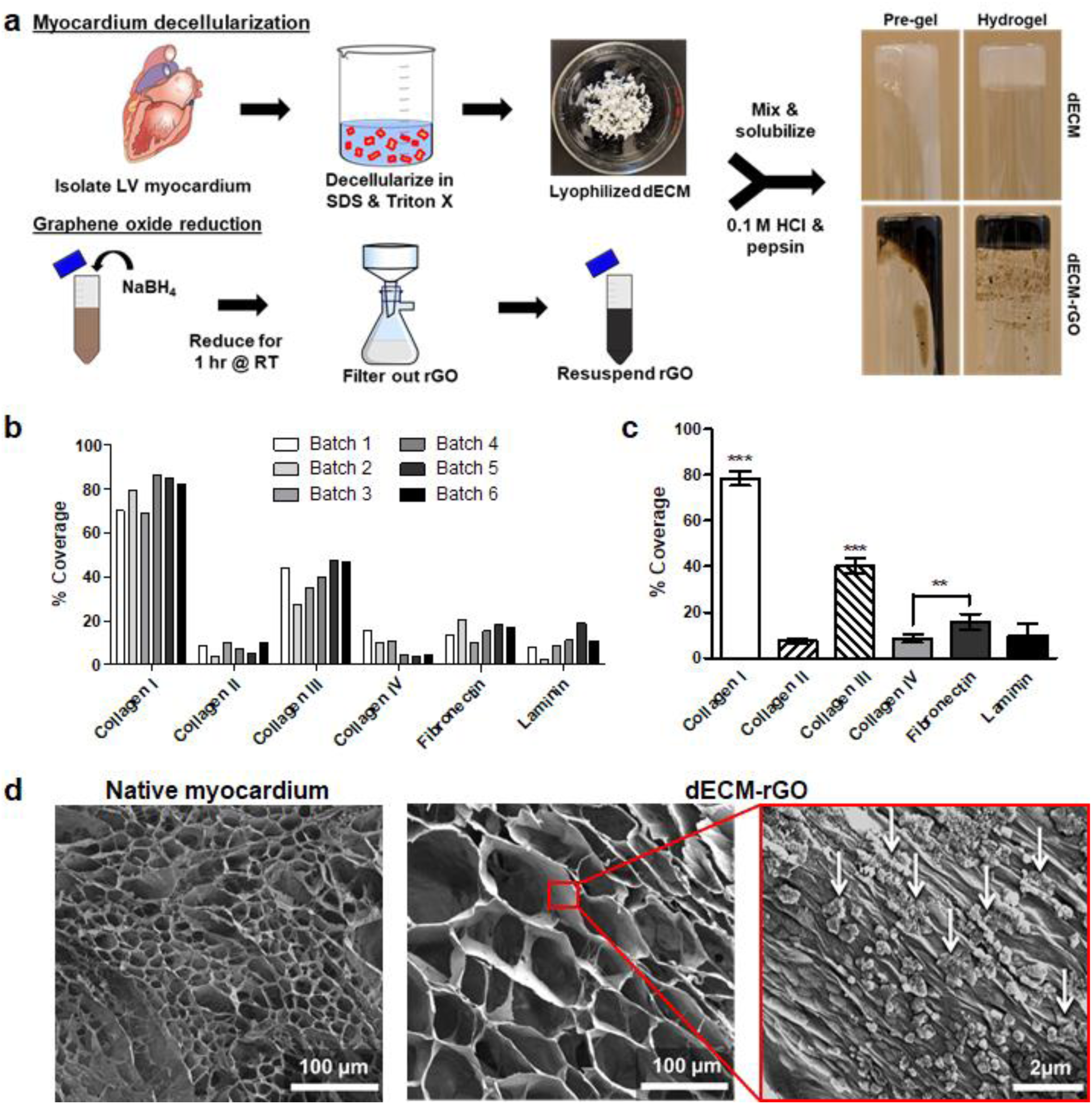
Materials synthesis and structural-biochemical characterization. (a) Composite hydrogels are synthesized using a parallel process. dECM (top) is produced by first isolating and mincing left ventricular myocardium from freshly-harvested porcine hearts. Minced tissue is decellularized using a combination of detergents before being lyophilized. rGO (bottom) is produced by reducing GO with NaBH4 for 1 hr before filtration and resuspension in dH2O. The two components are combined with HCl and pepsin to form a pre-gel solution that can then be formed into hydrogels by incubation at 37°C. (b) LC/MS analysis of dECM indicates that while some batch-to-batch variation occurs, (c) overall protein composition is relatively well maintained across produced batches. **p < 0.01, ***p < 0.001 (One-way ANOVA with a Tukey’s *post-hoc* test, n = 6). (d) SEM imaging of dECM-rGO hydrogels reveals a porous structure similar to that of decellularized myocardium. Deposition of rGO flakes (white arrows) on the pore walls can be observed.

Reduction of the graphene oxide (GO) was accomplished using a sodium borohydride (NaBH4)-mediated chemical reduction process in which NaBH4 removes oxygen groups from the GO lattice structure in a similar manner to hydrazine (N2H4), but without the use of a highly-toxic compound^20^. By adjusting the concentration of NaBH4 used in the reduction process, the degree of reduction of GO to rGO could be varied, and this was confirmed with the use of Raman spectroscopy (**Supplementary Fig. 1**). As NaBH4 concentration increased, the D:G peak intensity ratio increased, indicating a removal of lattice defects and a shift towards the restoration of a near-pristine graphene lattice structure.

Examination of the structure of composite dECM-rGO hydrogels with scanning electron microscopy (SEM) showed that the hydrogels possessed a porous network similar to that of decellularized native porcine myocardium (**Fig. 1d**). Closer examination of dECM-rGO pore walls revealed that nanoscale flakes of rGO were deposited on the walls, indicating that encapsulated cells would be able to interact closely with the rGO. It is unclear, however, whether these rGO aggregates penetrated through the walls such that adjacent pores would be connected or bridged by the electroconductive material.

The mechanical properties of dECM-rGO hydrogels were examined using compressive and rheological testing. Compressive modulus was found to increase as both a function of the degree of reduction of the rGO (**Fig. 2a**) and as a function of rGO content within the hydrogels (**Fig. 2b**). The average compressive moduli obtained with the hydrogels tested in this study ranged from 10.6 ± 0.3 kPa (dECM only) to 17.5 ± 0.5 kPa (dECM with 0.3% w/v 300 mM NaBH4-reduced rGO), which are comparable to the range for native myocardium^21, 22^. Hydrogel viscosity was also found to increase as rGO reduction (**Fig. 2c**) and content increased (**Fig. 2d**). Notably, all dECM-based hydrogel formulations exhibited shear-thinning properties, in which material viscosity decreased as shear rate increased. Storage (G’) and loss (G”) moduli were found to change in response to hydrogel composition in the same manner as before, with G’ and G” values increasing as rGO reduction and content increased (**Supplementary Fig. 2**).

**Figure 2.**
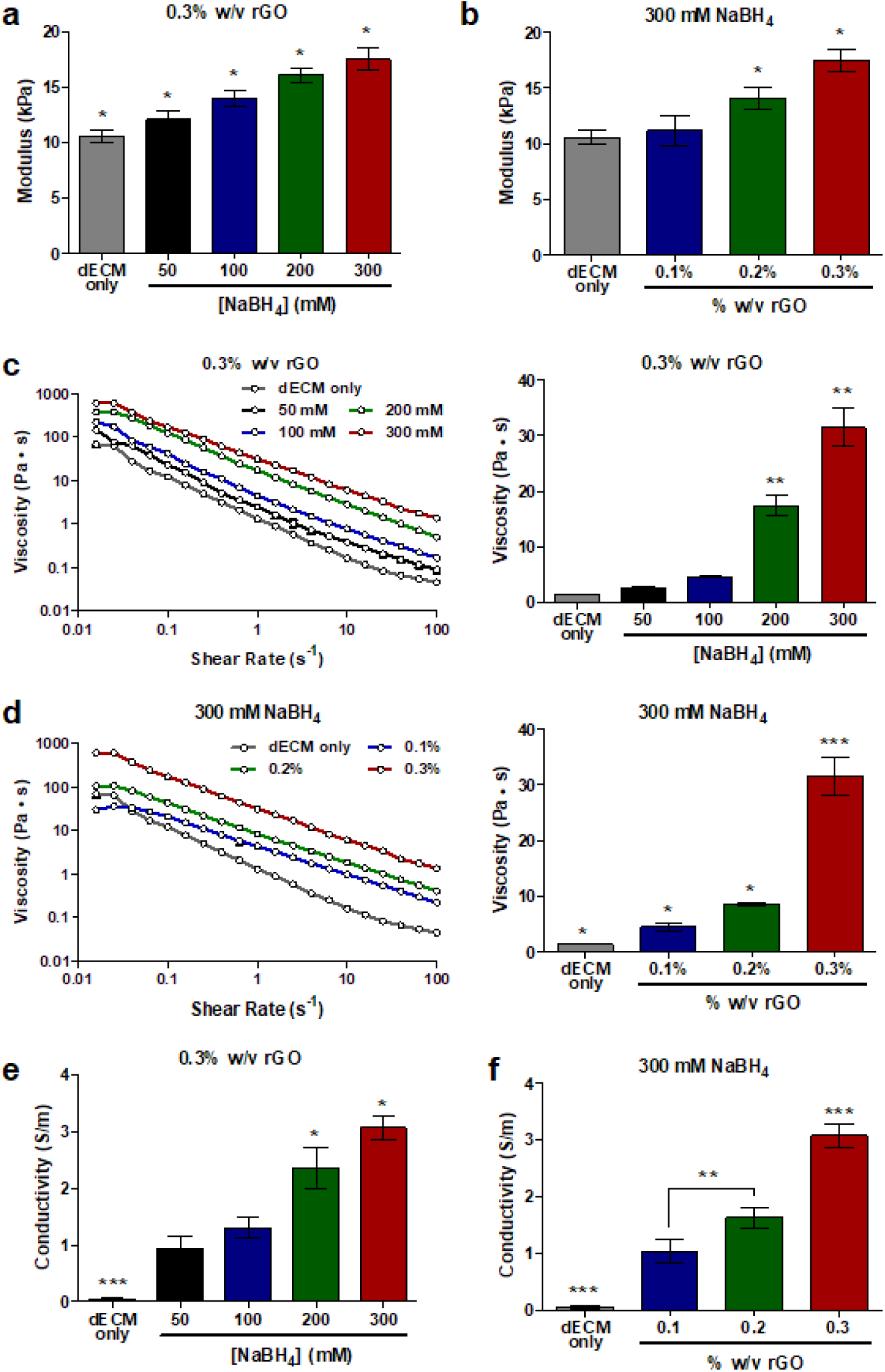
The mechanical and electrical properties of dECM-rGO is tunable. Compressive modulus of composite hydrogels increases as a function of both (a) the degree of reduction of rGO, and (b) the rGO content within the composite hydrogels when degree of reduction is held constant. (c, d) Similar trends can be observed in the viscosity of dECM-rGO pre-gel solutions, and these solutions also exhibit shear-thinning properties. Hydrogel conductivity also increases as a function of (e) degree of rGO reduction and (f) rGO content. *p < 0.05, **p < 0.01, ***p < 0.001 (one-way ANOVA with a Tukey’s *post-hoc* test, n = 6).

Dehydrated scaffolds were subjected to four-point probe measurements to determine the effect of rGO on material electroconductivity. dECM-only scaffolds were not highly conductive, with a measured average conductivity (0.0005 ± 0.0002 S/m). Increasing rGO reduction and content led to corresponding increases in hydrogel electroconductivity **(Fig. 2e, f**), with average hydrogel conductivities ranging from 0.93 ± 0.12 S/m to 3.07 ± 0.12 S/m. While these values are approximately 3-10 times greater than those reported for native myocardium^23, 24^, this could be beneficial for supporting a high degree of electrical signal propagation during the early stages of cell and tissue maturation when gap junctions have not yet been fully developed. In summary, it was demonstrated that *via* the nature and content of the incorporated rGO, the mechanical and electrical properties of the resultant dECM-rGO hydrogels could be tuned. It should be noted that the observed increases in stiffness due to rGO could not be decoupled from a corresponding increase in conductivity; therefore, in order to have a dECM-based hydrogel that could serve as a stiffness control in subsequent cell-based studies, dECM hydrogels crosslinked with transglutaminase (TG) were utilized. These dECM-TG hydrogels featured compressive moduli similar to that of the stiffest formulation of dECM-rGO (**Supplementary Fig. 3**), and thus fulfilled the requirement for a hydrogel that still possessed the biochemical cues of dECM, but without the enhanced electroconductivity imparted by rGO.

Cytotoxic effects from highly-conductive carbon lattice-based materials, such as graphene, graphene oxide, and carbon nanotubes, have been reported in literature^25, 26^. An assessment of the biocompatibility of dECM-rGO hydrogels, however, demonstrated that the presence of rGO had no long-term negative effects on cardiomyocyte and stromal cell viability (> 90% viable after 35 days), even at the highest concentrations used in this study (**Supplementary Fig. 4**).

### Enhanced engineered heart tissue contractile function and maturation mediated by dECM-rGO hydrogels

Three-dimensional engineered heart tissues (EHTs) were generated by casting a mixture of hiPSC-derived cardiomyocytes, human bone marrow-derived stromal cells, and pre-hydrogel solution around silicone posts that enabled *in situ* twitch force measurements *via* post deflection imaging (**Fig. 3a, Supplementary Videos 1 & 2**). In these studies, collagen I hydrogel was selected as a control scaffold material that was closest in approximation to the biochemical make-up of dECM and is commonly used to fabricate 3D cardiac tissues. Additionally, the stiffest and most electroconductive dECM-rGO composition (dECM with 0.3% w/v 300 mM NaBH4 rGO) was used, and as mentioned previously, dECM-TG served as a stiffness control. EHTs were cultured over a 35-day period, with their contractile performance analyzed every 7 days. While no difference in twitch force was found in the first 7 days, a significant increase in force output was detected in dECM-rGO tissues on Day 14 (**Fig. 3b**). Specifically, the dECM-rGO tissues produced twitch forces nearly twice that of the next closest group (23.61 ± 2.62 μN vs. 12.97 ± 3.00 μN, dECM-rGO vs. dECM-TG, respectively), and nearly five times that of the collagen tissues (23.61 ± 2.62 μN vs. 5.33 ± 2.36 μN). Noting this striking improvement in contractile function at Day 14, all subsequent experimental endpoints and data analyses were performed for Day 14 and Day 35 time points. dECM-rGO tissues continued to produce greater twitch forces compared to the other tissue groups over the entire culture period, with dECM-TG tissues consistently generating the next greatest force output, followed by dECM-only tissues, and lastly by collagen tissues. However, it was interesting to observe that at Day 28 and Day 35, it appeared that the advantage in force generation by dECM-rGO tissues was beginning to diminish, with dECM-TG and dECM tissues producing nearly identical twitch forces (approximately 45 μN). Meanwhile, force output of the collagen tissues had essentially plateaued by Day 28 to 28.99 ± 1.37 μN. Twitch power generated by EHTs followed a similar trend, with dECM-rGO tissues producing greater power at both Day 14 and Day 35 while no significant difference was found between dECM and dECM-TG tissues at the latter time point (**Supplementary Fig. 5**).

**Figure 3.**
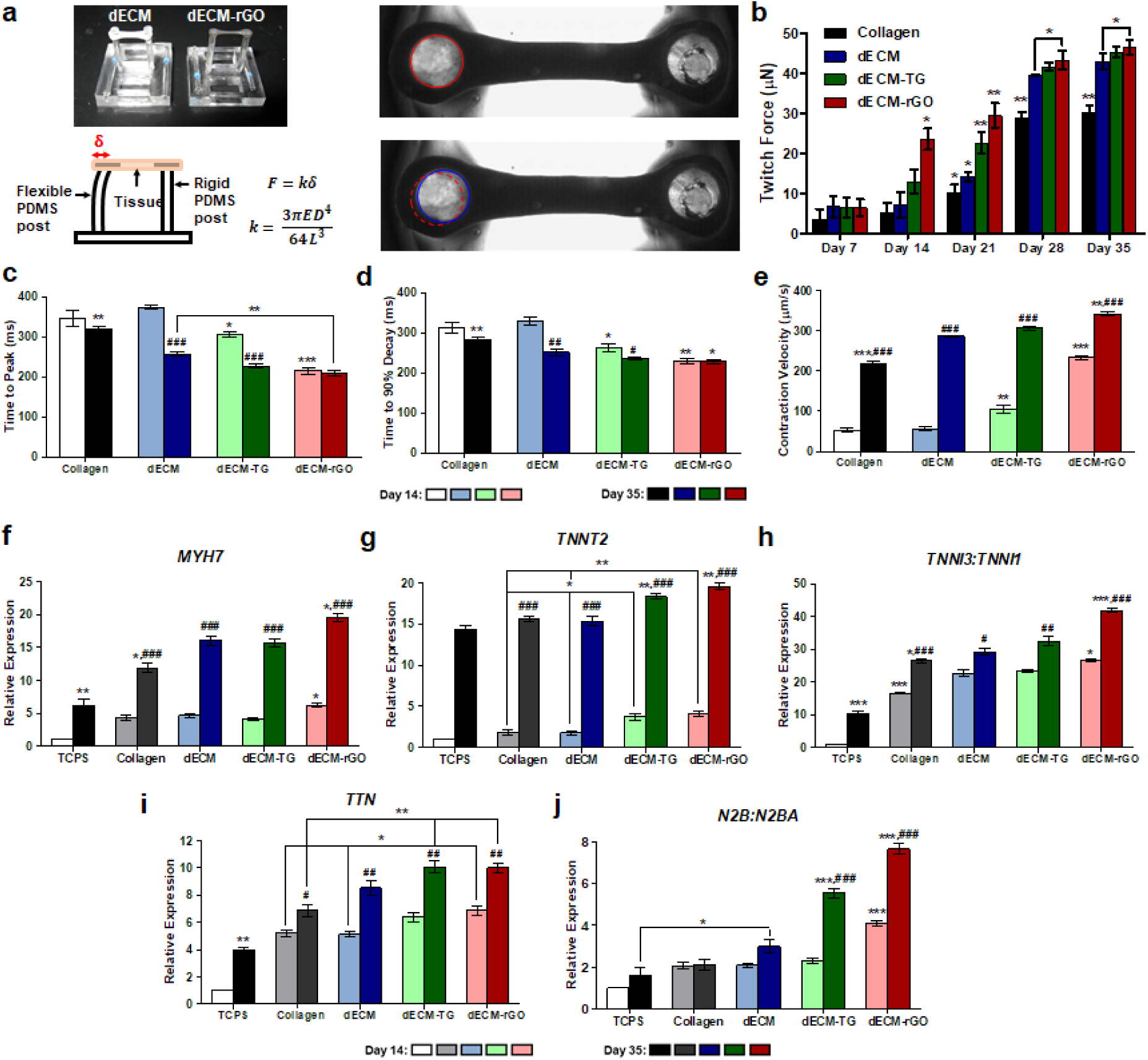
Contractile function is enhanced in dECM-rGO heart tissues. (a) Representative photographs of dECM and dECM-rGO tissues formed on the two-post EHT platform (top left) and schematic illustrating the beam theory that allows for quantitative analysis of contractile function using the EHT platform, where E = elastic modulus of PDMS, D = diameter of the post, L = length of the post, and δ = measured post deflection (bottom left). Snapshots of twitching EHTs (right), in which post deflection is indicated by the relative position of the post cap at two different time points (red and blue circles). (b) Twitch forces generated by dECM-rGO tissues is significantly greater than those of other tissues at all time points after Day 7, with the greatest difference observed on Day 14. Twitch forces appear to plateau after Day 28. p < 0.05, **p < 0.01 (one-way ANOVA with a Tukey’s *post-hoc* test, n = 12). (c) dECM-rGO tissues generate maximal twitch forces quicker than other groups at both Day 14 and Day 35, and (d) reach 90% relaxation quicker than other groups at both time points as well. (e) As such, dECM-rGO tissues feature significantly greater contraction velocities than tissues formed with other hydrogels. (f) RT-qPCR analysis of *MYH7* shows that expression levels in dECM-rGO tissues is greater than controls that show no significant difference between them at Day 14, and is still greater at Day 35. (g) Expression of *TNNT2* in dECM-TG and dECM-rGO tissues is greater at both time points, with no significant difference observed between collagen and dECM tissues. (h) Relative expression ratios of cardiac (*TNNI3*) and skeletal (*TNNI1*) isoforms of troponin I show a greater induced expression of the cardiac-specific gene in dECM-rGO tissues. (i) Overall expression of titin (*TTN*) is increased in dECM-rGO tissues on Day 14, but this increase is not statistically-significant from that of dECM-TG. On Day 35, both dECM-TG and dECM-rGO have increased expression of *TTN* relative to collagen, but this difference is not significant compared to dECM. (j) However, when looking at the expression ratios of *N2B* to *N2BA*, dECM-rGO tissues show a significant increase at Day 14, while both dECM-rGO and dECM-TG tissues have vastly larger expression levels relative to others at Day 35. *p < 0.05, **p < 0.01, ***p < 0.001 (hydrogel material comparison; one-way ANOVA with a Tukey’s *post-hoc* test, n = 12); ^#^p < 0.05, ^##^p < 0.01, ^###^p < 0.001 (time point comparison; Student’s *t*-test, n = 12).

In examining the contractile kinetics of the EHTs, it was found that not only did the dECM-rGO tissues outperform the rest in terms of twitch force generation, but were also able both to reach maximal force (**Fig. 3c**) and relax to 90% force (**Fig. 3d**) at a faster rate per contraction, thereby attaining significantly greater contraction velocities (**Fig. 3e**). Notably, there was no significant difference in times to peak and 90% relaxation between Days 14 and 35 for dECM-rGO tissues, while all other tissue groups displayed improvements in these metrics over time. Additionally, while the contraction kinetics of dECM and collagen tissues were comparable to each other at Day 14, by Day 35, dECM tissues were outperforming their collagen counterparts.

With the observed enhancement of EHT contractile performance due to dECM-rGO hydrogel scaffolds, the expression of several genes that code for cardiomyocyte contraction-mediating proteins was analyzed and compared to hiPSC-derived cardiomyocytes grown as monolayers on tissue culture polystyrene (TCPS) as controls. Expression of *MYH7*, which codes for β-myosin heavy chain (β-MHC), was significantly more elevated in dECM-rGO tissues on Day 14 (**Fig. 3f**). *MYH7* expression was significantly increased for all groups at Day 35, but expression levels in other tissues were still lower than those seen in the dECM-rGO group. Similar trends were observed for *TNNT2*, or cardiac troponin T (cTnT), although in this case both dECM-TG and dECM-rGO tissues had greater expression levels at both time points compared to the others (**Fig. 3g**). Upregulation of these genes corresponded well with the collected twitch force data, as cTnT and β-MHC regulate the force generation capacity of cardiomyocytes and are considered late-stage markers of cardiac development^27–29^. Troponin I is another critical component of the myocyte contractile machinery, and a comparison of troponin I isoforms can also shed light on the maturation state of cardiomyocytes, as cells with a fetal or neonatal phenotype will largely express *TNNI1* (slow skeletal TnI/ssTnI), while adult cardiomyocytes will exclusively express *TNNI3* (cardiac TnI/cTnI)^5, 30, 31^. Thus, the significantly higher ratio of expression of *TNNI3* to *TNNI1* in dECM-rGO tissues at both time points indicates that this hydrogel composition enhances the maturation of these tissues, which likely is an important driver of improved contractile function (**Fig. 3h**). Additionally, although there is no significant difference in *TNNI3*:*TNNI1* between dECM and dECM-TG tissues, both possess a greater amount of the cardiac isoform than the collagen and TCPS controls. At both time points, the expression levels of titin (*TTN*), the protein responsible for muscle elasticity, were comparable between dECM-TG and dECM-rGO, which were in turn greater than that in dECM and collagen (**Fig. 3i**). Furthermore, the ratio of the *N2B* to *N2BA* isoforms of titin was significantly higher in dECM-rGO tissues at both Day 14 and Day 35, another indication that the maturation state of these cardiomyocytes was improved by the composite hydrogel (**Fig. 3j**). Indeed, the increased presence of the N2B isoform in adult myocardium and resulting increase in passive stiffness is what allows for rapid ventricular recoil^32^, which correlates well with the observed decrease in relaxation time in dECM-rGO tissues.

Sarcomeres are the basic contractile unit of striated muscle and their structural properties directly impact force production^33^. As such, EHTs were sectioned and immunostained for sarcomeric α-actinin to assess how sarcomere development was influenced by scaffold material composition (**Fig. 4a**). Sarcomeres were shortest in collagen tissues (1.78 ± 0.01 μm) and progressively lengthened as the hydrogel scaffolds were more biochemically diverse, stiffer, and electroconductive, with the longest sarcomeres (2.11 ± 0.02 μm) found in dECM-rGO tissues (**Fig. 4b**). For comparison, sarcomeres in adult human cardiomyocytes are reported to be approximately 2.2 μm in length^34^, while immature cardiomyocytes cultured on standard 2D substrates typically have sarcomeres that are only about 1.5 – 1.65 μm long^35^. In addition to sarcomere length, the lateral alignment of adjacent sarcomeres plays an important role in contraction synchronicity and overall force output. Sarcomere register is typically assessed as a function of Z-band width, where cardiomyocytes capable of generating more twitch power feature wider Z-bands^36^. Similar to the trend observed with sarcomere length, Z-bands were shortest in collagen tissues (3.12 ± 0.12 μm) and widest in dECM-rGO tissues (3.92 ± 0.13 μm), although statistical difference was only found for the dECM-rGO group (**Fig. 4c**).

**Figure 4.**
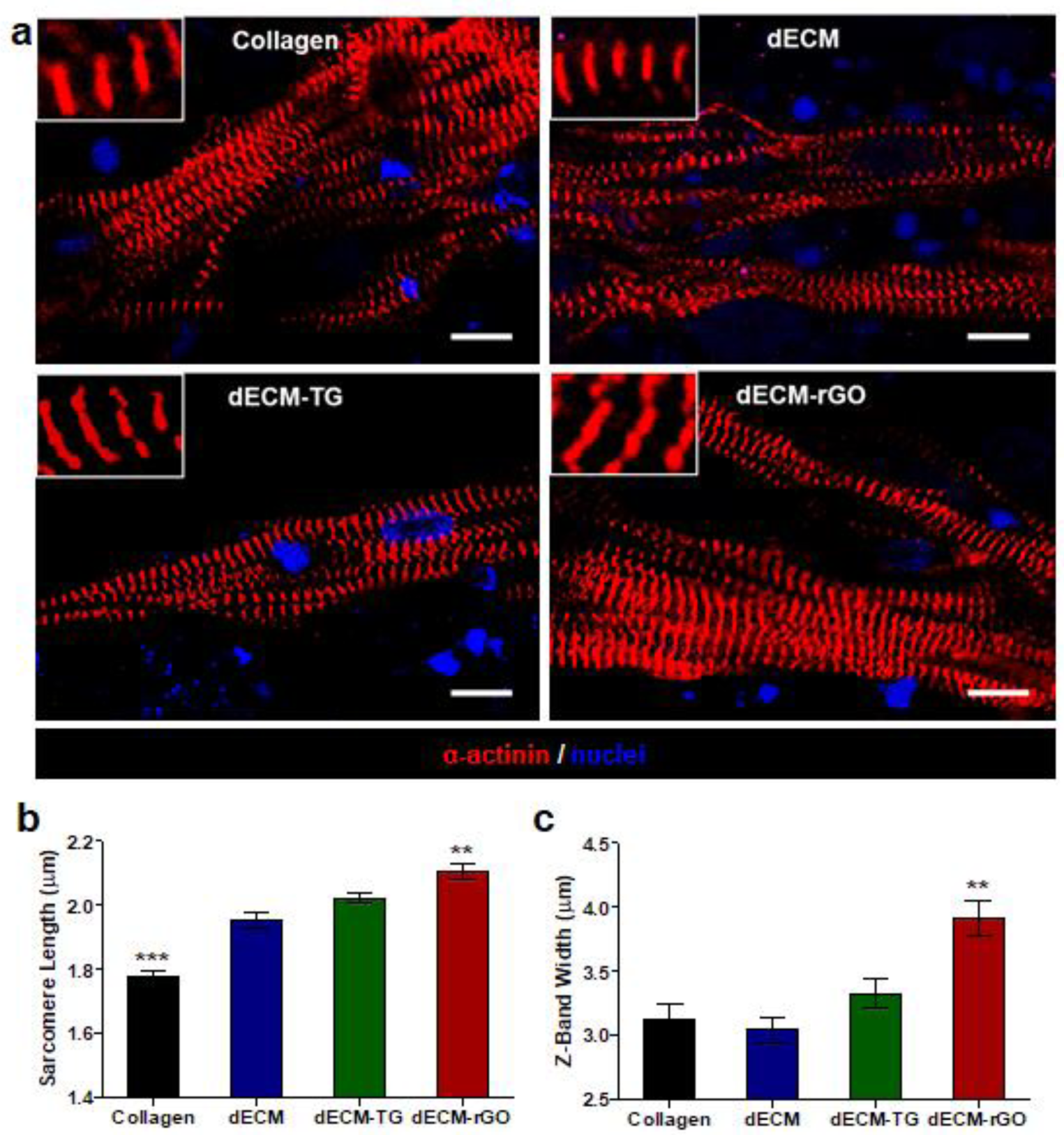
Sarcomere development in engineered heart tissues. (a) Representative fluorescent images of Day 35 tissues generated with the different hydrogels stained for sarcomeric α-actinin (red) and nuclei (blue). Scale bar = 10 μm. (b) Sarcomere length and (c) Z-band widths are increased in dECM-rGO tissues relative to other tissues, with sarcomere lengths approaching those observed in adult human cardiomyocytes. **p < 0.01, ***p < 0.001 (one-way ANOVA with a Tukey’s *post-hoc* test, n = 6).

### Enhanced EHT electrophysiological function mediated by dECM-rGO hydrogels

Calcium plays an important role in regulating the excitation-contraction coupling processes that are required for proper cardiac function. To determine the impact of scaffold material on the Ca^2+^-handling capabilities of EHTs, a Ca^2+^-sensing fluorescent dye was used to monitor changes in intracellular Ca^2+^ levels during contractions (**Fig. 5a**). At both Day 14 and Day 35, the normalized average change in fluorescence intensity was found to be significantly greater in dECM-rGO tissues versus its dECM counterparts, an indication of increased Ca^2+^ flux and thus a greater concentration of calcium present in these tissues each contraction cycle (**Fig. 5b**). Additionally, while Ca^2+^ transients increased over time for dECM-rGO tissues, transient levels remained relatively unchanged for dECM tissues. An examination of expression levels for ion channels involved in maintaining cytosolic calcium homeostasis, *CACNA1C* (L-type Ca^2+^ channel CaV1.2) and *ATP2A2* (Ca^2+^-ATPase; SERCA2), shows significantly elevated expression of both genes in dECM-rGO tissues compared to all other tissue types, with *ATP2A2* expression further increased on Day 35 (**Fig. 5c, d**).

**Figure 5.**
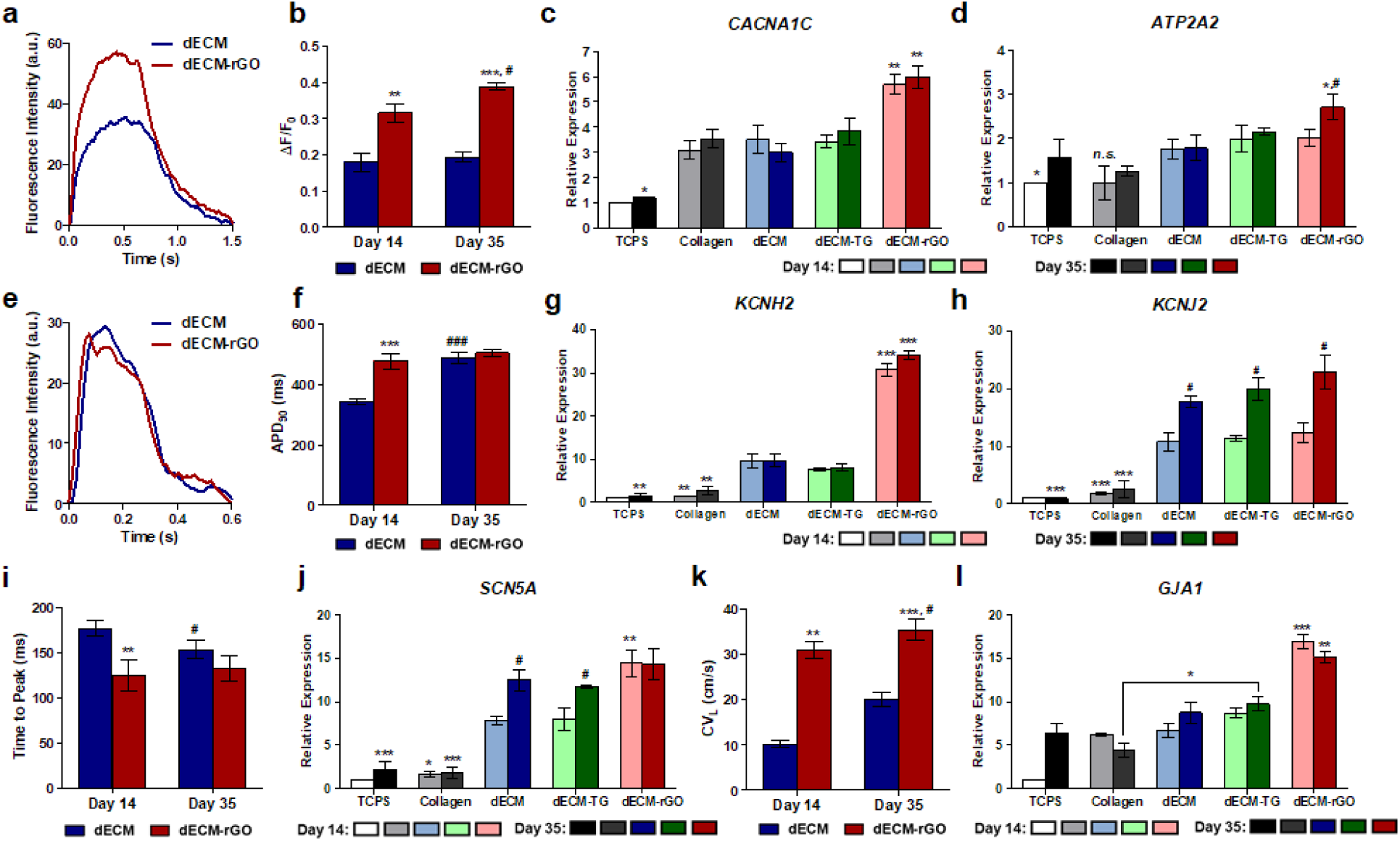
Electrophysiological function is also enhanced in dECM-rGO heart tissues. (a) Representative trace of fluorescence intensity obtained from live-tissue imaging of Fluo-4 AM at Day 35. (b) Change in fluorescence is significantly greater in dECM-rGO tissues at both Day 14 and Day 35, indicating greater Ca^2+^ flux across cardiomyocyte membranes. (c) RT-qPCR analysis of *CACNA1C* and (d) *ATP2A2* indicates that expression of these Ca^2+^ channel components is greater in dECM-rGO tissues, suggesting that calcium-handling capabilities are improved by the composite hydrogel scaffold. (e) Representative trace of fluorescence intensity obtained from live-tissue imaging of FluoVolt at Day 35. (f) Quantitative analysis of voltage dye traces shows that APD90, an analog for QT interval, is lengthened in dECM-rGO tissues at Day 14. By Day 35, there is no significant difference in APD90 values between both groups. (g) RT-qPCR analysis of *KCNH2* (hERG channel) expression shows a dramatic increase in dECM-rGO tissues across both time points. (h) Expression of *KCNJ2*, on the other hand, was comparable across all tissues engineered with dECM-based scaffolds although expression increased over time. (i) Time to peak depolarization is significantly reduced in dECM-rGO tissues compared to dECM counterparts. (j) Expression of *SCN5A* is significantly greater in dECM-rGO tissues at Day 14, but not on Day 35. (k) Longitudinal conduction velocity is greater in dECM-rGO tissues at both Day 14 and Day 35, and this corresponds with (l) a marked increase in *GJA1* (Cx43) expression in these tissues. *p < 0.05, **p < 0.01, ***p < 0.001 (hydrogel material comparison; one-way ANOVA with a Tukey’s *post-hoc* test, n = 12); ^#^p < 0.05 (time point comparison; Student’s *t*-test, n = 12).

To further characterize the electrophysiological function of these EHTs, a voltage-sensing fluorescent dye was used to investigate action potential behavior (**Fig. 5e**). Action potential duration at 90% repolarization (APD90), commonly used as an analog for QT interval, was observed to be significantly longer in dECM-rGO tissues (488 ± 21 ms) than in dECM tissues (343 ± 11 ms) at Day 14. At Day 35, however, both tissues had similar APD90 values at 477 ± 25 ms and 504 ± 12 ms, for dECM and dECM-rGO, respectively (**Fig. 5f**). For comparison, QT intervals in adult humans are generally reported to be in the 350 – 460 ms range^37^. In examining the expression of several K^+^ channels that contribute to the plateau phase of cardiomyocyte action potentials, it was found that *KCNH2* (human ether-a-go-go-related-gene; hERG) (**Fig. 5g**) and *KCNE1* (**Supplementary Fig. 6a**) were upregulated in dECM-rGO tissues by almost 3-fold at both Day 14 and Day 35, with no significant change in expression between the two time points. Meanwhile, expression of *KCNJ2* (KIR2.1) (Fig. 5h) and *KCNQ1* (**Supplementary Fig. 6b**) was not significantly different between tissues generated with dECM-based hydrogels; however, expression levels were much greater than that of the collagen and TCPS controls. Moreover, expression of these two genes was found to increase significantly over time for all three of the dECM-based hydrogel groups. The variations in the expression levels of these particular K^+^ channels (along with the increased *CACNA1C* exoression) may explain the observed trends in APD90. The significant upregulation of *KCNH2* and *KCNE1* in dECM-rGO tissues could account for its longer APD90 at Day 14. Then, as additional K^+^ channels such as *KCNJ2* and *KCNQ1* become more numerous in dECM tissues over time, APD90 values begin to equalize, as seen with the Day 35 analysis. In addition to an increase in APD90 mediated by dECM-rGO, the time to peak depolarization was significantly reduced at Day 14 in these tissues **(Fig. 5i**). Similar to the trend observed for APD90, there was no significant difference in time to peak depolarization between dECM and dECM-rGO tissues at Day 35. Expression of *SCN5A*, which codes for the voltage-gated Na^+^ channel NaV1.5 that is responsible for mediating membrane depolarization in cardiomyocytes, was found to be much higher in dECM-rGO tissues at Day 14, although no difference in expression was found amongst the three dECM-based tissues at Day 35 despite an overall upregulation across these groups (**Fig. 5j**). Like with the K^+^ channel expression patterns, this result likely explains the comparable depolarization rates between dECM and dECM-rGO tissues that were seen at the later time point.

Data collected from the voltage-sensing dye, FluoVolt, was also used to calculate the longitudinal conduction velocity (CVL) in EHTs. At both time points, conduction velocity was significantly greater in dECM-rGO tissues than in their dECM counterparts, with a peak CVL of 35.4 ± 2.3 cm/s recorded at Day 35 (**Fig. 5k**), which approaches the mean conduction velocity of isolated human left ventricular myocardium (49.8 ± 3.8 cm/s)^38^. Since electrical signal transmission between cardiomyocytes has been shown to be closely associated with connexin 43 (Cx43) gap junction formation^39^, the expression of the gene that codes for this protein was examined. Indeed, *GJA1* expression was significantly greater in dECM-rGO tissues, although no difference in expression levels was seen between the early and late time points in all tissues (**Fig. 5l**).

### Regulation of transcription factor expression by hydrogel composition

Transcription factors are known to play an important role in determining cell fate and tissue morphogenesis. In developing cardiac tissues, Nkx2.5 and GATA4 are highly expressed in the pre-cardiac mesoderm and in cardiac progenitor cells, and numerous gain/loss of function studies have shown that these two factors work in concert to drive the cardiomyocyte differentiation process^40–42^. RT-qPCR analysis of *NKX2.5* and *GATA4* expression in EHTs at Day 14 showed that although all four of the tested hydrogel scaffolds induced higher expression levels than what were seen in 2D cultured cells, only dECM-rGO tissues had a significantly greater expression level than collagen controls (**Fig. 6**). However, as EHTs became more mature over time, the expression of both genes saw a significant decrease. This reduction in expression was found to be closely tied to the type of scaffold material used; as the microenvironmental cues became more complex, the decrease in expression became greater. In fact, TCPS actually elicited an increase in expression in cardiomyocytes. These results serve to illustrate the impact on maturation rate that is elicited by 3D dECM-based hydrogels, and by dECM-rGO in particular.

**Figure 6.**
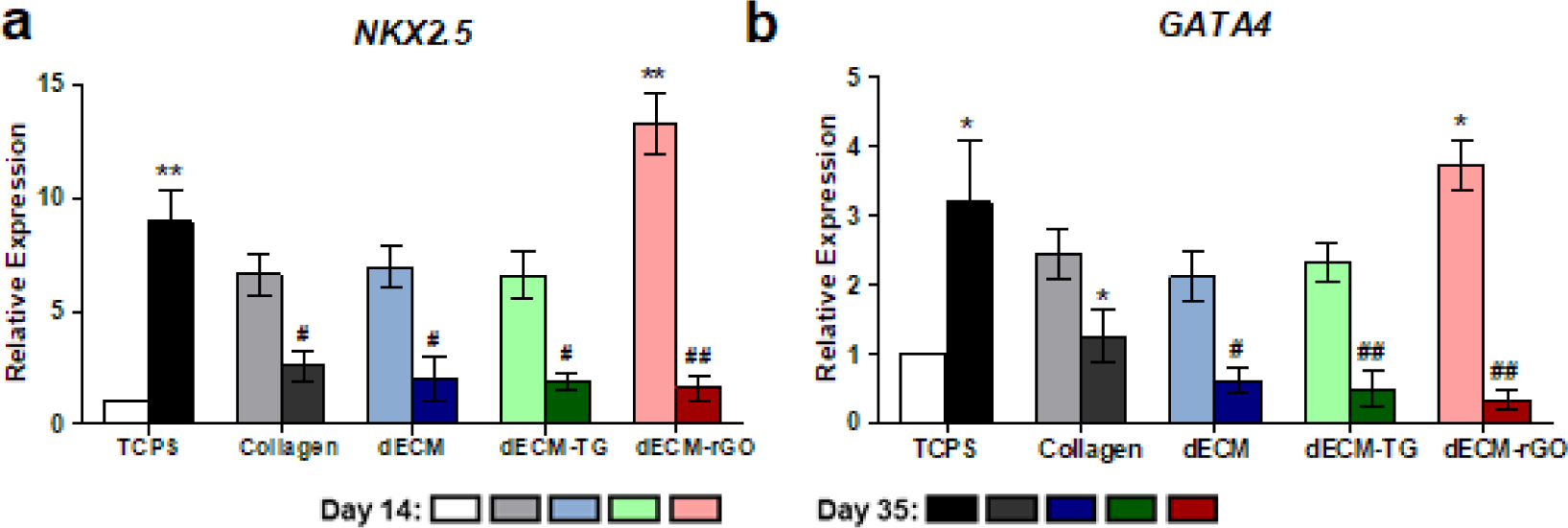
Hydrogel-mediated transcription factor expression over time. (a) dECM-rGO elicited a significant increase in expression of *NKX2.5* at Day 14, and all other 3D tissues had higher expression levels than TCPS controls. By Day 35, expression was comparable between all tissues. (b) Similar trends in relative expression over time were observed for *GATA4*, where dECM-rGO tissues had high levels of expression at Day 14 but had reduced expression by Day 35. *p < 0.05, **p < 0.01 (hydrogel material comparison; one-way ANOVA with a Tukey’s *post-hoc* test, n = 12); ^#^p < 0.05, ^##^p < 0.01 (time point comparison; Student’s *t*-test, n = 12).

### Bioprinting a microphysiological drug screening platform with dECM-rGO bioink

As noted previously, dECM and dECM-rGO hydrogels exhibited shear-thinning behavior in response to increases in strain rate. This type of viscoelastic property makes these hydrogels well-suited to applications such as bioprinting where it is desirable to have a material that can be easily extruded, while maintaining the capacity to quickly reform and hold a structure post-extrusion. Using a dual-extruder bioprinter, a multi-material design was developed in which hydrogels could be utilized as a bioink for generating 3D cardiac tissues in a high-throughput manner while taking advantage of the beneficial effects on tissue maturation that dECM-rGO could offer (**Fig. 7a**). In this design, pairs of vertical posts comprised of polycaprolactone (PCL) were printed to serve as anchor points for tissues and to induce anisotropic cellular alignment, while Pluronic F-127 was used to print temporary wells around these posts to aid in tissue formation during the first 24 hrs.

**Figure 7.**
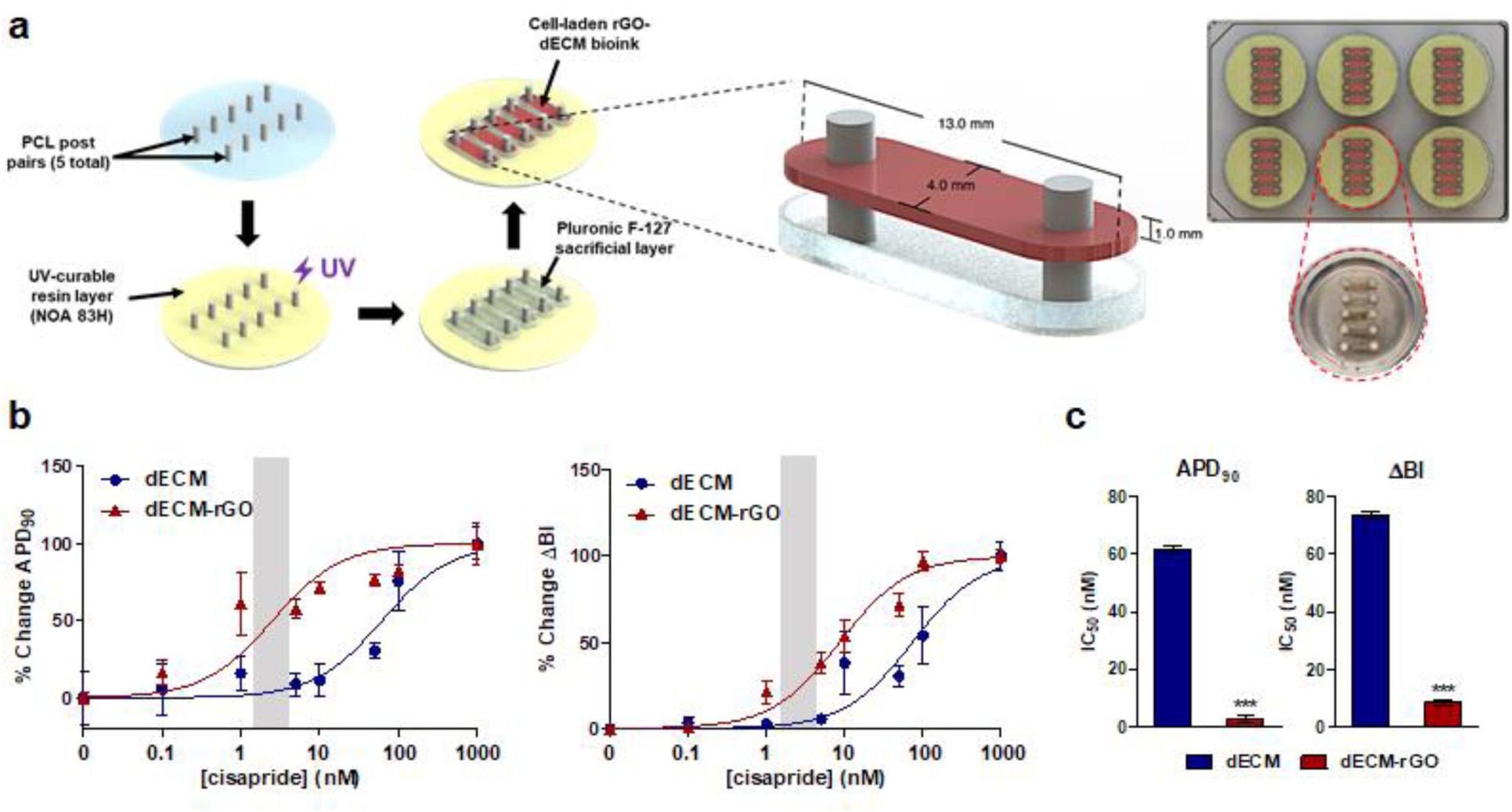
Bioprinted cardiac tissues can serve as a high-throughput drug-screening platform. (a) Schematic representation of the developed bioprinted platform. Post pairs comprised of PCL are printed and anchored into place with a thin layer of photocured NOA 83H. A temporary well of Pluronic F-127 is printed around each post pair before cell-laden dECM or dECM-rGO bioink is printed into each well. As an example, a 5 tissue-per-well in a 6 well plate setup is demonstrated. (b) Dose-response curves for percent change in APD90 and ΔBI in response to varying concentrations of cisapride in tissues cultured for 14 days. (c) Quantification of IC50 values derived from both metrics show that dECM-rGO tissues respond in a clinically relevant manner compared to dECM tissues at this time point. Gray boxes indicate reported ETPC range for cisapride. ***p < 0.001 (Student’s *t*-test, n = 10).

Using this platform, cardiac tissues were printed with dECM and dECM-rGO bioinks and cultured for 14 and 35 days before being subjected to a dose response study with cisapride. Cisapride is a histamine receptor antagonist later discovered to be a potent hERG channel blocker that induces QT interval prolongation and potentially lethal Torsade de Pointes (TdP). This arrhythmogenic property was not detected using conventional preclinical screening assays, and the arrhythmogenic complications led to its withdrawal from market^43–45^. Fluorescent imaging of FluoVolt dye was once again used to assess cisapride’s impact on tissue APD90 length and beat interval regularity across a large range of concentrations. At Day 14, it was found that dECM and dECM-rGO tissues responded significantly differently to increasing cisapride concentrations (**Fig. 7b**). While both tissue types exhibited APD90 prolongation and an increase in ΔBI, the IC50 values differed by an order of magnitude (**Fig. 7c**). dECM-rGO tissues responded to cisapride at clinically relevant concentrations, with IC50 values (APD90: 2.5 ± 1.3 nM; ΔBI: 8.6 ± 1.2 nM) that were very similar to its effective therapeutic plasma concentration (ETPC) of 2.6 – 4.9 nM^46^. On the other hand, dECM tissues had IC50 values that were much higher at 61.8 ± 1.4 nM and 73.4 ± 1.4 nM for APD90 and ΔBI, respectively. Surprisingly, there was no significant difference in drug response between the dECM and dECM-rGO tissues at Day 35, with best-fit dose response curves for percent change in APD90 and ΔBI identical between the two tissue types (**Supplementary Fig. 7a**). However, IC50 values for both dECM (APD90: 2.0 ± 1.3 nM; ΔBI: 7.3 ± 1.2 nM) and dECM-rGO tissues (APD90: 1.8 ± 1.4 nM; ΔBI: 3.1 ± 1.3 nM) were representative of native cardiac tissue response when exposed to cisapride (**Supplementary Fig. 7b**).

## Discussion

The results of this study demonstrate the ability to generate a library of bioactive and electroactive hydrogels through the doping of dECM with rGO, and the capability of these hybrid hydrogels to improve the functional phenotype of engineered cardiac tissues such that they can produce physiological responses to external stimuli. The electrical and mechanical properties of dECM-rGO hydrogels were modulated by the reduction of GO to a more graphene-like sp^2^ hybridized lattice structure that imparts high conductivity and mechanical strength^47^, and this phenomenon has been observed in other instances where the addition of graphene or rGO to polymer matrices also resulted in corresponding increases in modulus and conductivity^48–51^. While the use of pristine graphene would have likely resulted in hydrogels with even greater electroconductivities and stiffnesses, the relatively hydrophobic nature of graphene compared to rGO does not lend itself well to homogenous dispersion within aqueous matrices, risking tissue-wide inconsistencies in cellular response. Furthermore, residual oxygen-containing groups in rGO could induce the spontaneous adsorption of proteins and thereby act as depots for biomolecules that contribute to cell differentiation and development^52^. Although transglutaminase-crosslinked dECM was only used as a stiffness control in this study, it is plausible that the modulus of dECM-rGO hydrogels could be further increased with the addition of transglutaminase, thereby providing yet another degree of tunability to the hybrid material. This could prove to be an immensely useful tool for certain applications, such as the modeling of fibrotic and hypertrophic ventricular myocardium, in which a stiffer cell microenvironment is desired^53^.

In terms of twitch force production and contraction kinetics, the performance of dECM-only and collagen tissues were comparable at the early time point, and this was also reflected in the expression profiles of the selected genes. However, as time went on, the dECM appeared to confer a distinct advantage on inducing maturation, with notable gains in functional output and the expression of genes encoding titin and cTnI. dECM-mediated improvements in electrophysiological function were observed to occur even sooner in culture, with the greatest effect seen on K^+^ and Na^+^ channel gene expression. These effects can be attributed to the fact that although dECM hydrogels are a homogenization of many intra-tissue niches and the decellularization process slightly alters native tissue protein composition, compared to single component biomaterials like collagen, dECM-based materials are unique in that they are still capable of presenting a cell microenvironment with a wider spectrum of tissue-specific biochemical cues that are essential for supporting proper development. Indeed, similar findings have been reported in which dECM scaffolds have been found to exert enhanced differentiation and development on a variety of tissue types *in vitro*^18, 54–56^, as well as pro-regenerative effects on skeletal muscle^57–59^ and myocardium^60, 61^ *in vivo*.

Like most other naturally-derived scaffold hydrogels, however, there is a modulus mismatch between dECM and native tissue. The proper tuning of matrix stiffness has proven to be a critical factor in successfully engineering cardiac tissues, as prior studies have found that the development of cardiomyocyte contractile function and striation is significantly improved in microenvironments that closely approximate myocardial stiffnesses^22, 36, 62^. This is somewhat unsurprising, as there is a large body of evidence that implicates the role of integrins in translating biomechanical signals to determine cell fate and that myofibril assembly starts at these integrin adhesion sites^63–65^. The results presented here are consistent with those findings, as the stiffer dECM-TG and dECM-rGO hydrogels generated tissues that produced greater twitch forces and velocities, and an increased expression of *TNNT2*, *TNNI3*, *TTN*, and *N2B*. Sarcomere formation was also improved by a corresponding increase in matrix stiffness. Surprisingly, the relative expressions of the various ion channels and *GJA1* in dECM and dECM-TG tissues were statistically comparable, which suggests that matrix stiffness appears to have a limited effect on regulating this subset of electrophysiological development. Further examination of integrin-mediated signaling pathways is clearly needed; however, a study conducted with neonatal rat ventricular cardiomyocytes showed that even though substrates with biomimetic stiffnesses induced increased action potential duration and calcium flux, L-type Ca^2+^ channel expression in these cells was unchanged^66^.

Based on the stark improvement of EHT contractile and electrophysiological function and increased expression of associated developmental markers within these tissues, it is evident that rGO exerts a profound effect on multiple aspects of hiPSC-derived cardiomyocyte maturation. While there is an abundance of examples in which electroconductive scaffolds have been shown to enhance the maturation and function of cardiomyocytes^13, 48, 67–70^, the mechanisms and signaling pathways mediated by these types of materials have yet to be fully elucidated. In almost all reported cases, however, the most significant improvements in performance were in parameters related to electrical coupling, such as Cx43/*GJA1* expression, which has a direct correlation to increased conduction velocities. This implies that although electroconductivity still plays a role in upregulating other cardiac-specific proteins and transcription factors, it appears that the largest impact is on promoting electrophysiological development of these cells. The recent discovery that cells cultured on graphene exhibited an enrichment of the cholesterol content in their membranes raises interesting questions on what downstream effects this would potentially regulate^71^. For example, since membrane cholesterol has been shown to promote G protein-coupled receptor (GPCR) activity^72^, an increase in cholesterol could result in a corresponding increase in the activity of GPCRs, such as sphingosine 1-phosphate receptor-1, that contribute to normal cardiac development^73^. It has also been found that gap junctions like Cx43 and KV channels like hERG localize to cholesterol-rich membrane domains, and that alteration of this membrane lipid environment can modulate the function of these proteins^74–76^. Furthermore, an increase in plasma membrane cholesterol content has been shown to increase Ca^2+^ flux through membrane calcium channels in a number of different cell types, including cardiomyocytes^77–79^.

Thus, with the synergistically beneficial combination of biochemical, mechanical, and electroconductive cues offered by dECM-rGO hydrogels, a more pronounced maturation response in cardiomyocytes was observed than in tissues generated with only collagen or dECM. Improvements in some aspects of maturation were also evident when compared to other 3D human stem cell-based cardiac tissue engineering approaches that used non-conductive hydrogel scaffolds with a limited spectrum of tissue-specific biochemical cues^6, 80, 81^. However, it should be noted that the expression of some genes associated with the cardiomyocyte contractile machinery, namely *MYH7* and *TNNT2*, was found to be relatively lower than those in which engineered tissues underwent cyclic physical conditioning^7, 8^, suggesting that if dECM-rGO scaffolds were to be utilized in similar bioreactor systems, further gains in contractile phenotype maturation could be achieved.

A significant limitation of current preclinical screening techniques for drug-induced cardiotoxicity is their inability to provide accurate predictability, leading to incidences of false positives or negatives, increases in drug development costs, and unnecessary risk to patients. The reliance on animal models and non-human cell lines has contributed to this problem due to mismatches in pharmacokinetics and physiology between species^82^, so a shift towards the utilization of 3D humanized *in vitro* tissue models is a promising first step. However, not only do these approaches suffer from the aforementioned issues with phenotypic immaturity, but they are also hampered by low throughput and fabrication variability. The emergence of bioprinting and additive manufacturing as tissue fabrication methods offers a potential solution to these drawbacks as these techniques allow for the precise deposition and patterning of multiple biomaterials in a reproducible and automated fashion^83^. In the design developed in this study each post pair and well was a single defined unit, and this imparted increased modularity and versatility compared to more conventional grid designs^18, 84^ as it could be adapted for a variety of layouts to increase or decrease desired experimental throughput. For example, while a 5 tissue per well of a 6-well plate format was used, the same design could be used to print a single tissue per well in a 24-well or 48-well plate. When used with dECM-rGO bioinks, bioprinting was therefore able to produce larger numbers of more mature cardiac tissues in less time and in a format that still allowed for conventional image-based analysis of changes in electrophysiological function. While current hardware limitations do not allow for a level of throughput commensurate with existing systems such as microelectrode arrays (MEAs)^85^, the ability to utilize 3D tissues instead of 2D monolayers is still an advantageous aspect of this approach.

Based on the vast difference in expression of the hERG channel coding gene *KCNH2* between dECM and dECM-rGO tissues, it is perhaps unsurprising that even after only 14 days of culture, cisapride induced abnormal APD90 prolongation and beat intervals in dECM-rGO tissues at physiological concentrations. In comparison to previously reported drug screening platforms based on hiPSC-derived cardiomyocytes^85, 86^, whole animal models^87^, and *in silico* simulations^88^, our platform was able to produce IC50 values for cisapride or similar hERG channel blockers that were within the same order of magnitude as ETPC. What was surprising, however, was the indistinguishable difference in response of the dECM and dECM-rGO tissues to cisapride after 35 days of culture despite there still being a significant gap in *KCNH2* expression at this time point. This result could indicate that as time progressed, dECM scaffolds were able to induce further development of other ion channels that were not examined in this study and that these channels contributed to an overall maturation state that allowed these tissues to behave physiologically to this particular drug. In fact, other studies have suggested that the onset of TdP cannot be solely explained by the blockade of a single current^89^. However, when combined with the collected twitch force data, the results of this cisapride assay could also serve as further evidence of the role dECM-rGO hydrogels play in accelerating the maturation of engineered cardiac tissues to the point that even though the developmental stage has not yet reached a fully adult-like phenotype, these tissues are still mature enough to produce useful data in drug screening and disease modeling applications. To the end user, this could potentially provide significant advantages to cost and throughput, as tissues would only need to be maintained for a shorter duration of time before use.

In summary, the findings of this study illustrate the synergistic impact that the microenvironmental cues present in dECM-rGO hydrogels had on promoting the functional maturation of engineered cardiac microphysiological systems. Not only did this biomaterial induce significant improvements in several metrics of cardiac function, these developmental gains were translated into clinically-relevant physiological responses suitable for downstream applications. Furthermore, the tunability of these hybrid hydrogel’s properties and demonstrated applicability as a bioink for 3D bioprinting enable it to be a versatile tool for answering multiple biological questions. The ease in which two materials with largely disparate characteristics could be combined into a single composite demonstrates the potential for other adjuncts that could be incorporated into these hydrogels, such as soluble or tethered factors or other synthetic nanomaterials that could be used to further enhance overall material functionality.

## Methods

### Tissue decellularization

Myocardial decellularized extracellular matrix (dECM) was obtained by an adaptation of well-established methods detailed in the literature^18, 90^. In brief, left ventricular myocardium was isolated from heats collected from freshly-slaughtered adult pigs and sectioned into small slices approximately 1 mm in thickness. Slices underwent cell lysis and removal in a solution of 1% w/v sodium dodecyl sulfate (SDS; Fisher Scientific) for 36 h, with an additional wash in 1% w/v Triton-X100 (Sigma-Aldrich) for 1 hour to complete cellular material removal. This was followed by washes in phosphate buffered saline (PBS) over 3 days, with the wash solution changed every day to remove excess detergent and cell debris. dECM was sterilized with 0.1% w/v peracetic acid in 4% w/v ethanol, followed by an additional wash in PBS overnight. Afterwards, the dECM was lyophilized and stored at 4°C until used.

### Characterization of dECM biochemical composition

Dry weights of lyophilized dECM were determined and the samples were digested at a concentration of 5 mg/mL in a solution containing 5 M urea, 2 M thiourea, 50 mM DTT and 0.1% SDS in PBS with constant stirring at 4°C for 48 h. Afterwards, the samples were sonicated on ice (Branson Digital Sonifier, 20 s pulses, 30% amplitude), and protein was precipitated with acetone and analyzed *via* liquid chromatography tandem mass spectrometry (LC-MS/MS). The six most abundant ECM components for each developmental age were identified from spectrum count data.

### Graphene oxide reduction and reaction validation

Reduced graphene oxide (rGO) was chemically reduced from commercially-available graphene oxide solution (GO; Graphene Laboratories) using NaBH4 (Sigma-Aldrich), as previously described^20^. Briefly, GO flakes with a lateral flake size of 90 nm – 200 nm in a 1 mg/mL stock solution were mixed with varying concentrations of NaBH4 (50 mM – 300 mM) for 1 hour at room temperature. The resulting rGO was then vacuum filtered through mixed cellulose ester membranes (MilliporeSigma) with a 0.1 μm pore size. Filtered rGO was then removed and resuspended in dH2O at 10 mg/mL using a sonication bath. Prior to resuspension of filtered rGO, samples were collected for reduction process validation and analysis with a confocal Raman microscope (Renishaw InVia). The relative intensity of the D (1350 cm^−1^) and G (1600 cm^−1^) peaks were then measured from the collected spectra of each sample.

### Extracellular matrix solubilization and hydrogel formation

dECM-based pre-hydrogel solutions were generated by mixing lyophilized dECM and pepsin at a 10:1 mass ratio in 0.1 M HCl. rGO was also added at this stage for dECM-rGO hydrogels. A 2% w/v dECM solution was used for all experiments in this study while the final concentration of the rGO component was varied from 0.1% – 0.3% w/v. Mixtures were stirred constantly until the dECM was completely solubilized and had formed a homogenous solution, at which point the pH was then brought up to 7.4 with 10 M NaOH. Finally, 10% of the final volume of 10X PBS or 10X RPMI 1640 (Sigma-Aldrich) was added to restore isotonic balance. Hydrogel pre-gel solutions were stored at 4°C until use. Hydrogels were then formed by incubating pre-gel solutions at 37°C for at least 12 h. The same solubilization and gelation method was used to generate control hydrogels composed of bovine collagen type I (Advanced BioMatrix). For generating dECM-TG hydrogels for stiffness controls, a 5% w/v solution of transglutaminase (TG; Modernist Pantry) was added to the dECM solution immediately prior to incubation at 37°C.

### Microstructural imaging with scanning electron microscopy

dECM-rGO hydrogels and decellularized myocardial sheets underwent critical-point drying. Samples were then sputter-coated with a thin layer of Au/Pd alloy before imaging in a FEI Sirion XL30 scanning electron microscope at a 5 kV accelerating voltage.

### Characterization of hydrogel mechanical and electrical properties

Hydrogel conductivity was quantified using a four-point probe (Four Dimensions Model 280SI). Thin films were deposited onto a glass coverslip and dehydrated. Probe measurements were taken at 20 different points across the sample and averaged. Compressive moduli of hydrated crosslinked hydrogels were measured using an Instron compressive testing system, with the composite hydrogel samples compressed at a rate of 10 mm/min until failure. Rheological measurements were taken with a rheometer (TA Instruments Discovery HR-2,) using two different protocols. Viscosity values were obtained by conducting a shear rate sweep of 0.1 – 100, with uncrosslinked pre-gel solution held at 25°C. G’ and G” values were obtained by conducting a frequency sweep of 0.1 – 100 rad/s at 1% strain, with crosslinked hydrogels held at 25°C. For both protocols, tested samples first underwent a 10 s pre-shear at 0.1 rad/s.

### Human induced pluripotent stem cell culture and differentiation

UC 3-4 urine-derived human induced pluripotent stem cells (hiPSCs)^3, 91^ were cultured and differentiated using an adaptation of a previously published method of modulating Wnt/β-catenin signaling with small molecules^92^. In brief, hiPSCs were maintained on 1:60 diluted Matrigel-coated plates in mTeSR1 medium (STEMCELL Technologies) prior to induction. Induction is achieved by culturing the cells in RPMI 1640 medium (Invitrogen) supplemented with B-27 without insulin (Invitrogen) and 10 μM CHIR-99021 (Selleck Chemicals) for 18 hrs, after which the medium is replaced with fresh RPMI/B-27 without insulin for 48 hrs. The medium is then replaced with RPMI/B-27 without insulin supplemented with 5 μM IWP-4 (REPROCELL) and cells are cultured for 48 hrs, after which the medium is once again replaced with fresh RPMI/B-27 without insulin for 48 hrs. Finally, medium is replaced with RPMI/B-27 with insulin (Invitrogen), and cultures are fed with fresh medium every other day. Spontaneously beating cells are typically observed 10-12 days after induction. Differentiation runs that produced hiPSC-cardiomyocyte (hiPSC-CM) purity of >90% as measured by flow cytometry after immunofluorescent staining with a human cardiac troponin T (cTnT) mouse monoclonal antibody (Thermo Fisher Scientific) were used for all experiments (**Supplementary Fig. 8**).

### Stromal cell source and culture

HS-27A human bone marrow-derived stromal cells (ATCC) were cultured and maintained in high-glucose DMEM (Invitrogen) supplemented with 10% fetal bovine serum until used to generate tissues.

### Assessment of biocompatibility

dECM only and dECM-rGO scaffolds containing 0.3% w/v rGO of varying degrees of reduction were seeded with hiPSC-CMs and HS27A stromal cells and cultured for 35 days, after which samples were stained with a live/dead fluorescent assay (Invitrogen) following the protocol provided by the manufacturer and imaged using a widefield epi-fluorescence microscope (Nikon Instruments Eclipse Ti). Live cells appeared as green (calcein-AM excitation/emission λ: 488/515 nm), while dead cells appeared as red (ethidium homodimer-1 excitation/emission λ: 570/602 nm). Quantitative analysis of captured images was performed using ImageJ (National Institutes of Health).

### Engineered heart tissue platform fabrication

The design and fabrication of the hardware utilized for both generating the engineered heart tissues (EHTs) and measuring forces *in situ* has been previously described^93, 94^. In brief, post pair arrays were fabricated by pouring uncured polydimethylsiloxane (PDMS; Sylgard 184, Dow Chemical) at a 10:1 base to curing agent ratio into an acrylic mold. A glass capillary tube was inserted into one post of each pair to impart rigidity to that post. Molds were then allowed to de-gas prior to being placed into a 65°C oven overnight to cure the PDMS. Cured post pair arrays were then removed from the mold. Arrays were fabricated in sets of six pairs, with each post 12.5 mm long and 1.5 mm in diameter and featuring a cap structure 0.5 mm thick and 2 mm in diameter to assist with tissue attachment. Post-to-post spacing was 8 mm. Wells for tissue casting were fabricated by pouring approximately 1 mL of PDMS into the wells of a 24-well culture plate before inserting custom 3D printed molds that would form rectangular wells that were 12 mm × 4 mm × 4mm (length × width × depth). After curing the PDMS overnight, molds were removed. All fabricated hardware was sterilized with 70% v/v ethanol and UV light prior to use.

### Generating engineered heart tissue constructs

hiPSC-CMs and HS-27A stromal cells were mixed in hydrogel pre-gel solutions at cell densities of 20 × 10^6^ cells/mL and 5 × 10^6^ cells/mL, respectively. 70 μL of this cell-hydrogel mixture was then pipetted into each casted PDMS well, after which PDMS post arrays were positioned upside-down into the wells, ensuring that each post tip was properly immersed in solution. The 24-well plate was then placed into an incubator at 37°C and 5% CO2 for approximately 60 min, after which the hydrogels had crosslinked and 1 mL of RPMI/B-27 with insulin (supplemented with Y-27632 dihydrochloride ROCK inhibitor and 1% penicillin/streptomycin) was added to each well. After 24 hrs, the media was replaced with fresh RPMI/B-27 with insulin and media changes were conducted every other day until tissue collection for endpoint analyses. Approximately 1 week after casting, tissues typically would have compacted sufficiently that post arrays with attached tissues could be removed from PDMS wells transferred into standard 24-well plates, at which point 2 mL of media per well was used for each media change.

### *In-situ* tissue function measurements and analysis

Prior to live-tissue imaging, tissues were transferred into a custom-built 24-well plate containing carbon electrodes with the wells filled with warmed Tyrode’s buffer with 1.8 mM Ca^2+^, and the tissues were allowed to stabilize for approximately 30 mins. The plate was then connected to a square-wave electrical stimulator (Grass Instruments S88X) and placed in a heated chamber mounted onto a widefield epi-fluorescence microscope. Videos of at least 5 contractions in duration were recorded using a monochrome CMOS camera (Mightex SMN-B050-U) with a board lens (The Imaging Source, TBL 8.4-2 C5MP) at 66 frames per second (FPS), with the tissues paced at 1.5 Hz and with a 5 V square wave amplitude. Image field of view included the entire tissue and the flexible and rigid posts. Using a custom MATLAB script, flexible post deflection was tracked across the duration of each video, which allowed for the calculation of twitch force, time to peak force, time to 90% force decay, contraction velocity, and twitch power^95^. In these calculations, slender-beam theory was applied, where the spring constant of the PDMS post, k, was determined by the Young’s modulus of PDMS (∼3.8 MPa) and the diameter and length of the post itself.

Intracellular Ca^2+^ flux and cardiac tissue depolarization characteristics were assessed using Fluo-4 AM (Invitrogen) calcium indicator and FluoVolt (Invitrogen) potentiometric dyes, respectively. Engineered heart tissues were washed with warm PBS before being incubated in medium containing either 5 μM of Fluo-4 AM or a 1000X dilution of stock FluoVolt for 1 hr at 37°C, after which tissues were washed twice with warm Tyrode’s buffer and placed in fresh Tyrode’s. Tissues were then imaged using standard FITC settings on a widefield epi-fluorescence microscope with a high-speed color CMOS camera (Hamamatsu ORCA-Flash4.0), with videos taken at 66 FPS and 2X magnification with a 0.7X coupler between the objective and camera. Tissues for calcium imaging were paced at 0.67 Hz with field stimulation generated by electrodes placed at each end of the tissue, and fluorescence intensity over time was quantified using NIS Elements software (Nikon) and used to calculate ΔF/F0 values. For action potential imaging, tissues were paced at 1.5 Hz and fluorescence intensity from the FluoVolt dye was again quantified using NIS Elements. APD90 and time to peak values were then calculated using a custom MATLAB script that removed motion artifacts from the analyses^94^. Longitudinal conduction velocity (CVL) was calculated by measuring the time delay in the onset of depolarization at two regions of interest that were at opposite ends of the tissue’s long axis and were a known distance apart.

### Gene expression analysis

The relative expression levels of selected genes were obtained using reverse-transcription quantitative polymerase chain reaction (RT-qPCR) analyses. After 14 and 35 days of culture, tissues were gently removed from posts and each tissue was placed into an Eppendorf tube containing a 100 μL solution of 2 mg/mL proteinase K in PBS. Tubes were then heated at 56°C for 10 mins before 350 μL of lysis buffer was added. The mixture was vortexed until clear and 250 μL of ethanol was added to the tubes. RNA was then subsequently isolated from this solution by using the RNeasy Plus RNA extraction kit (QIAGEN) according to the manufacturer’s protocol. cDNA was then synthesized from the isolated RNA with a reverse transcription kit (iScript Reverse Transcription Supermix, Bio-Rad). Quantity and purity of collected RNA and synthesized cDNA was determined by 260/280 nm absorbance. Synthesized cDNA was then analyzed with qPCR using SYBR green (Bio-Rad) as the reporter and primers used as provided from the manufacturer (Bio-Rad). Amplicon context sequences for these primers are provided in **Supplementary Table** 1, and primer sequences for *N2B* and *N2BA* are provided in **Supplementary Table 2**. Relative gene expression was calculated using the comparative Ct method, where *GAPDH* was designated as the housekeeping gene and age-matched hiPSC-derived cardiomyocytes cultured on tissue culture polystyrene (TCPS) with the same ratio of HS27A stromal cells as used in the EHTs were used as controls. Each sample was run in triplicate for each gene.

### Immunofluorescent imaging of sarcomere development

EHTs were cultured for 35 days and fixed in 4% paraformaldehyde in PBS overnight at 4°C while the tissues were still mounted on PDMS posts. Fixed tissues were then washed in fresh PBS and placed into a 20% sucrose solution overnight at 4°C. Tissues were then gently removed from the posts and frozen in optimal cutting temperature (OCT, Tissue-Tek) compound and sectioned into 10 μm slices for storage. Sections were initially blocked with a solution of PBS with 5% goat serum and 0.2% Triton X-100 for 1 hour at room temperature. Afterwards, sections were incubated with primary antibodies for α-actinin (1:200 dilution; Sigma) overnight at 4°C. Sections were washed with PBS and incubated with secondary fluorescent antibodies (1:400 dilution; Invitrogen) for 3 hrs at room temperature, with another series of PBS washes afterwards. Mounting media containing DAPI (Vectashield, Vector Laboratories) was then applied along with a coverslip, and stained sections were then stored at 4°C in the dark until imaging. Tissue sections were imaged using a confocal microscope (Nikon Instruments A1R) at 60X magnification with oil immersion. Analysis of sarcomere lengths and Z-band widths was conducted using ImageJ software, where a minimum of 10 cells per field of view over 5 distinct fields of view at single Z planes per sample were analyzed.

### Cardiac tissue bioprinting

Tissues and support structures were bioprinted using a commercially-available dual-extruder pneumatic bioprinter (Allevi BioBot 1). Five pairs of polycaprolactone (PCL) posts 1 mm in diameter, 5 mm tall, and 10 mm apart were printed in each well of a standard 6-well plate. An approximately 2 mm thick layer of NOA 83H resin (Norland Optical Adhesives) was then deposited onto the bottom of each well and cured by a 30 min exposure to UV light. Plates containing anchored posts were then sterilized by immersion in 70% v/v ethanol and subsequent UV light exposure. Sacrificial wells 12 mm × 5 mm × 1 mm (length × width × depth) were then printed around each post pair using 40% w/v Pluronic F-127, followed by approximately 50 μL of cell-laden hydrogel bioinks with the same cell densities as used in the EHT platform (20 × 10^6^ hiPSC-derived cardiomyocytes/mL and 5 × 10^6^ HS27A stromal cells/mL). Completed plates were then incubated at 37°C and 5% CO2 for approximately 60 min, after which 5 mL of RPMI/B-27 with insulin (supplemented with Y-27632 dihydrochloride ROCK inhibitor and 1% penicillin/streptomycin) was added to each well. After 24 hrs, the media was replaced with fresh RPMI/B-27 with insulin and media changes were conducted every other day until tissues were used for drug screening. Optimized printing parameters for each component of the design is provided in **Supplementary Table 3**.

### Assessment of cardiotoxicity with cisapride

Cardiac tissues were bioprinted using dECM and dECM-rGO bioinks and cultured for 14 and 35 days. At these time points, tissues were washed with warm PBS before being incubated in medium containing a 1000X dilution of stock FluoVolt for 1 hr at 37°C, after which tissues were washed twice with warm Tyrode’s buffer and incubated in Tyrode’s containing 0.01, 0.1, 1, 5, 10, 50, 100, and 1000 nM cisapride (Sigma-Aldrich) dissolved in dimethyl sulfoxide (DMSO) for 45 mins. A group of tissues were also incubated in Tyrode’s containing no cisapride, but an equivalent volume of DMSO (1 μL DMSO/mL Tyrode’s). Functional response of these tissues to administered drug was assessed using FluoVolt dye in conjunction with live-tissue imaging as described above. Change in beat interval (ΔBI) was determined by measuring that duration of elapsed time between individual beats (the beat interval) and calculating the difference in this interval from one beat to the next. APD90 values were calculated as described above. Best-fit dose response curves were calculated using GraphPad Prism software. Drug concentration values were first transformed to a log scale and APD90 or ΔBI values were normalized such that the lowest value was set at 0% and the largest value at 100%. Finally, the normalized data was fit to a non-linear dose response curve using a least squares fit.

### Statistical analyses

Unless otherwise noted, all quantitative data is presented as means ± standard error with a minimum of n = 6. One-way ANOVA with a Tukey’s *post-hoc* test was used to analyze data sets that included more than two experimental groups, while a Student’s *t*-test was used to compare data sets looking at only two variables. In all presented analyses, p < 0.05 was considered significant and n is defined as a separate tissue.

## Acknowledgements

This work was supported by National Institutes of Health grants R01 HL135143, R21 EB020132, and UG3 EB028094 (to D.-H.K.), TL1 TR002318 (to J.H.T.), and F32 HL126332 (to A.L.), National Science Foundation grant CMMI-1661730 (to N.J.S.), and American Heart Association fellowship 16PRE30760018 (to J.H.T.). Part of this work was conducted at the Washington Nanofabrication Facility and Molecular Analysis Facility, which are National Nanotechnology Coordinated Infrastructure (NNCI) sites at the University of Washington supported in part by the National Science Foundation (grants ECC-1542101, NNCI-1542101, 1337840 and 0335765), the University of Washington, the Molecular Engineering & Sciences Institute, the Clean Energy Institute, the National Institutes of Health, the Washington Research Foundation, the M. J. Murdock Charitable Trust, Altatech, ClassOne Technology, GCE Market, Google and SPTS. The content of this report is solely the responsibility of the authors and does not necessarily represent the official views of the National Institutes of Health. The authors would like to thank John Foster for his device fabrication and bioprinting assistance, Prof. Buddy Ratner for the use of his laboratory’s compression testing equipment, Amrita Basu and Prof. Alshakim Nelson for their rheological analysis expertise, Dr. Lil Pabon for her input and assistance with stem cell differentiation optimization, and Dr. Alec Smith for his insight into the electrophysiological and drug-response behavior of our engineered tissues.

