## Supplementary Information for "Functional Maturation of Human iPSC-based Cardiac Microphysiological Systems with Tunable Electroconductive Decellularized Extracellular Matrices"

### Supplemental Material

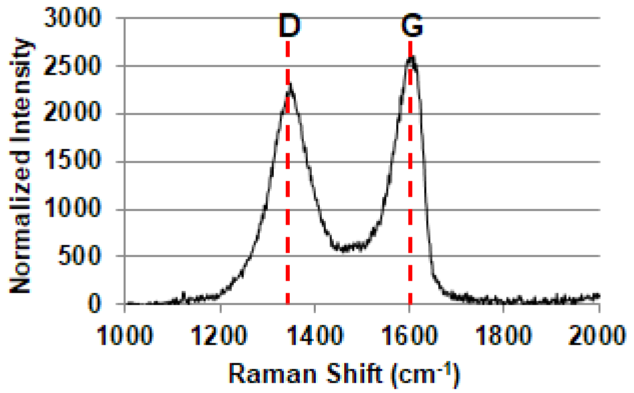


| [$NaBH_4$] (mM) | $I_D/I_G$ |
|---|---|
| 0 (neat GO) | 0.729 ± 0.031 |
| 50 | 0.817 ± 0.027 |
| 100 | 0.888 ± 0.019 |
| 150 | 0.867± 0.056 |
| 200 | 0.972 ± 0.035 |
| 250 | 1.082 ± 0.042 |
| 300 | 1.232 ± 0.061 |

**Supplementary Figure 1. Validation of graphene oxide reduction process.** Representative Raman spectra of GO illustrating the presence of the D and G peaks. Increasing the degree of reduction *via* $NaBH_4$ concentration leads to a corresponding increase in D peak intensity ($I_D$) relative to G peak intensity ($I_G$).

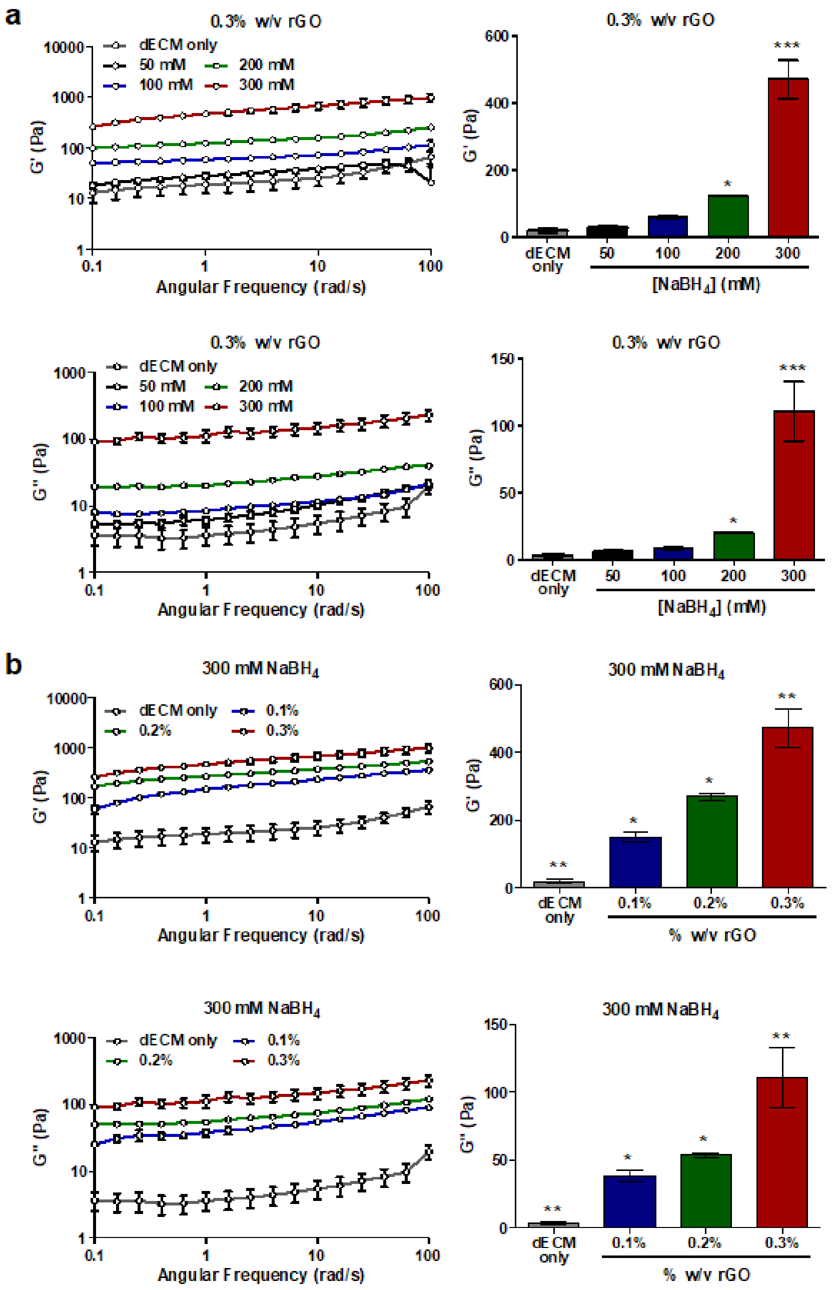


**Supplementary Figure 2. Tunable storage and loss modulus of dECM-rGO hydrogels.** The storage (G') and loss (G") moduli of dECM-rGO hydrogels increase with corresponding increases in (a) degree of rGO reduction and (b) overall concentration of rGO present in the dECM matrix. *$p < 0.05$, **$p < 0.01$, ***$p < 0.001$ (one-way ANOVA with a Tukey's *post-hoc* test, n = 6).

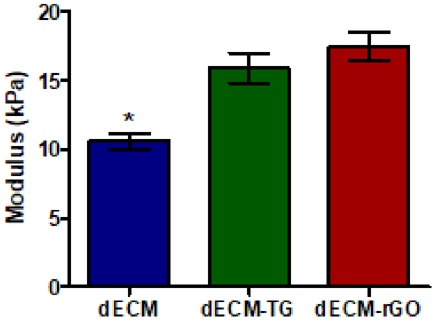


**Supplementary Figure 3. The compressive modulus of dECM can be increased with transglutaminase.** dECM hydrogels that were further crosslinked with transglutaminase (dECM-TG) possessed compressive moduli comparable to that of the stiffest dECM-rGO hydrogels (0.3% rGO reduced with 300 mM $NaBH_4$). *$p < 0.05$ (one-way ANOVA with a Tukey's *post-hoc* test, n = 6).

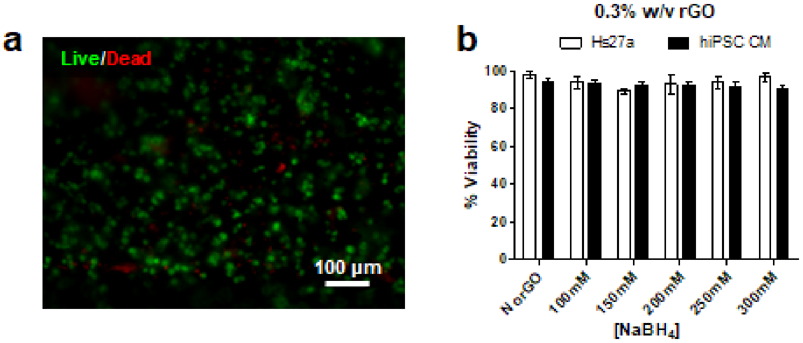


**Supplementary Figure 4. dECM-rGO hydrogel biocompatibility.** (a) Representative live/dead fluorescent image of a cardiomyocyte and stromal cell mixture cultured on dECM-rGO after 35 days. Live cells are labeled green and dead cells are labeled red. (b) High viability of both cell types is maintained even on hydrogels with high rGO content and degree of reduction.

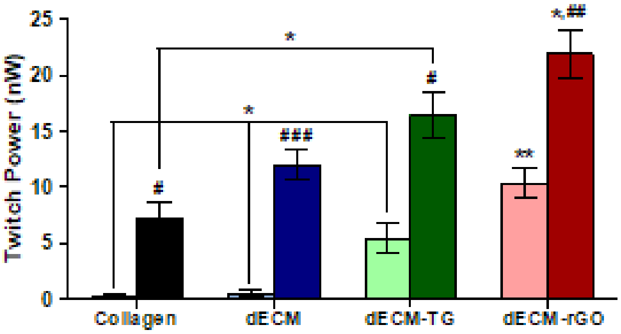


**Supplementary Figure 5. Twitch power generation by EHTs.** Twitch power generated by dECM-rGO tissues is significantly greater than controls at both time points. No significant difference is observed between dECM and dECM-TG tissues at Day 35. *$p < 0.05$, **$p < 0.01$ (hydrogel material comparison; one-way ANOVA with a Tukey's *post-hoc* test, n = 12); $^{\#}p < 0.05$, $^{\#\#}p < 0.01$, $^{\#\#\#}p < 0.001$ (time point comparison; Student's *t*-test, n = 12).

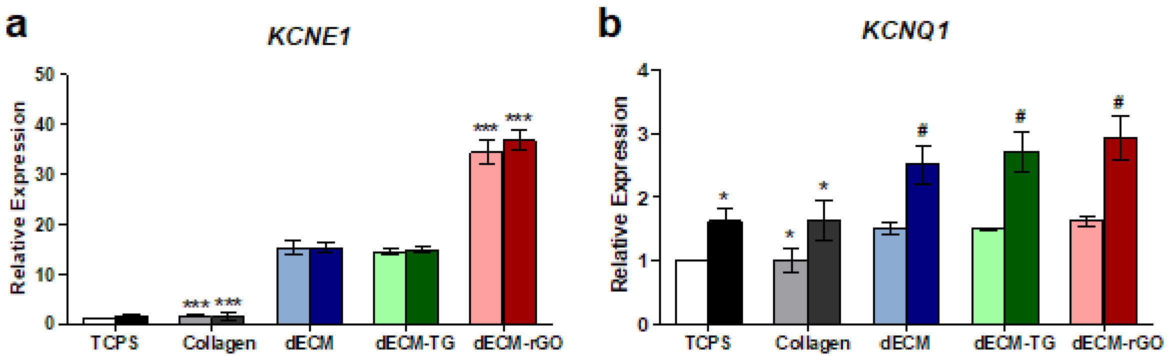


**Supplementary Figure 6. RT-qPCR analysis of additional $K^+$ ion channels.** (a) Expression of *KCNE1* was significantly greater in dECM-rGO tissues at both time points although expression levels did not appreciably change over time. (b) Expression of *KCNQ1* in all tissues generated with dECM-derived scaffolds was comparable to each other but still greater than in collagen at Day 14. Expression increased for all groups over time, but again significant differences were not found between dECM derivatives which still had greater levels of expression compared to collagen and TCPS. *$p < 0.05$, ***$p < 0.001$ (hydrogel material comparison; one-way ANOVA with a Tukey's *post-hoc* test, n = 12); $^{\#}p < 0.05$ (time point comparison; Student's *t*-test, n = 12).

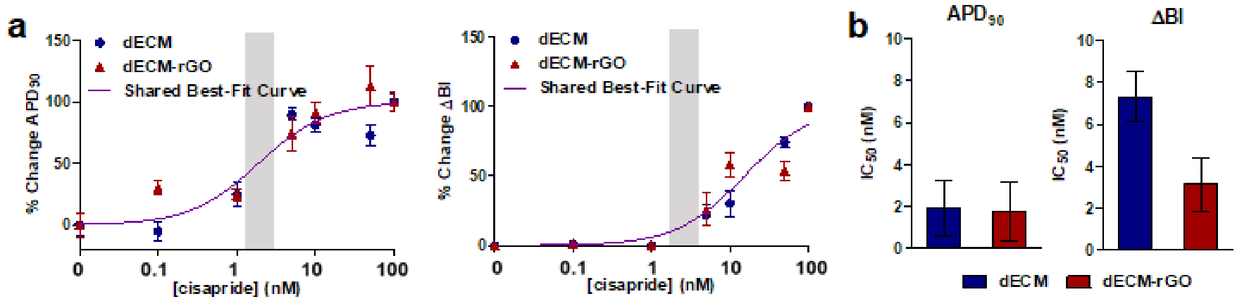


**Supplementary Figure 7. Bioprinted tissues at Day 35 do not show a differential response to cisapride.** (a) Bioprinted tissues cultured for 35 days showed no significant difference in their dose-response curves for percent change in $APD_{90}$ and ΔBI, and (b) this is reflected in the comparable $IC_{50}$ values for both metrics. Gray boxes indicate reported ETPC range for cisapride.

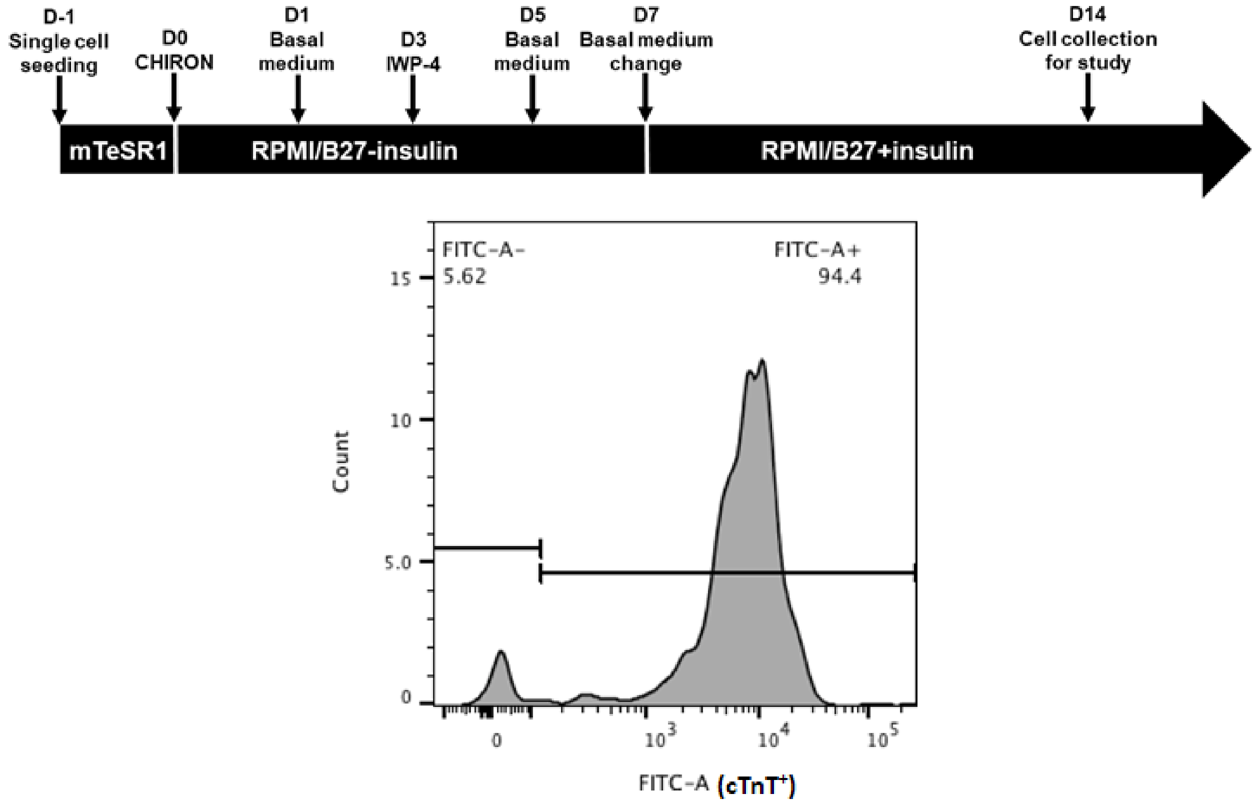


**Supplementary Figure 8. hiPSC cardiomyocyte differentiation scheme.** UC 3-4 hiPSCs were differentiated into cardiomyocytes using small molecule-mediated Wnt/β-catenin signaling modulating. Beating cardiomyocytes were typically observed 10 – 12 days after induction with CHIR-99021. A representative dataset from flow cytometry of cells positive for cardiac troponin T (cTnT) illustrates the high population purity (94.4%) that is typically achieved using this differentiation method.

**Supplementary Table 1.** Amplicon context sequences and lengths for primers used in RT-qPCR analyses.

| Gene | Amplicon Context Sequence | Amplicon Length (bp) |
|---|---|---|
| *MYH7* | 14:23891477-23892840 | 113 |
| *TNNT2* | 1:201330439-201331128 | 117 |
| *TNNI3* | 19:55665477-55666186 | 146 |
| *TNNI1* | 1:201379465-201380537 | 122 |
| *TTN* | 2:179665307-179666964 | 173 |
| *CACNA1C* | 12:2690767-2690902 | 106 |
| *ATP2A2* | 12:110765750-110770464 | 106 |
| *KCNH2* | 7:150649624-150649717 | 64 |
| *KCNJ2* | 17:68170965-68171100 | 106 |
| *KCNE1* | 21:35820801-35820975 | 145 |
| *KCNQ1* | 11:2799218-2869045 | 69 |
| *SCN5A* | 3:38614052-38620958 | 100 |
| *GJA1* | 6:121756921-121768079 | 130 |

**Supplementary Table 2.** Primer sequences for *N2B* and *N2BA*.

| Gene | Forward | Reverse |
|---|---|---|
| *N2B* | 5'-CCAATGAGTATGGCAGTGTCA-3' | 5'-TACGTTCCGGAAGTAATTTGC-3' |
| *N2BA* | 5'-CAGCAGAACTCAGAATCGA-3' | 5'-ATCAAAGGACACTTCACACTC-3' |

**Supplementary Table 3.** Printing parameters optimized for bioprinting microphysiological systems using dECM-based bioinks and a two-post passive tension design with a sacrificial support well

| Material | Pressure (psi) | Temp. (°C) | Needle Gauge |
|---|---|---|---|
| **PCL** | 100 | 100 | 30 |
| **Pluronic F-127** | 75-85 | 25 | 30 |
| **Bioink** | 3-5 | 25 | 25 |
| **NOA 83H** | 1-3 | 25 | 25 |
